# Adipocyte differentiation of 3T3-L1 cells under TAF, TDF and INSTIs selective challenge: an in vitro model

**DOI:** 10.1101/2022.07.16.500298

**Authors:** A. Perna, M.A. Carleo, S. Mascolo, A. Guida, M. Contieri, C Sellitto, E. Hay, P. De Blasiis, A. Lucariello, G. Guerra, A. Baldi, A. De Luca, P. Maggi, V. Esposito

**Affiliations:** Department of Medicine and Health Sciences “Vincenzo Tiberio”, University of Molise, Campobasso, Italy; Infectious diseases and Gender Medicine Unit, Cotugno Hospital, AO dei Colli, Naples, Italy; Department of Mental and Physical Health and Preventive Medicine, Section of Human Anatomy, University of Campania “Luigi Vanvitelli”, Naples, Italy; Department of Sport Sciences and Wellness, University of Naples “Parthenope”, Naples, Italy; Department of Environmental, Biological and Pharmaceutical Sciences and Technologies, University of Campania “Luigi Vanvitelli”, Caserta, Italy

**Keywords:** Weight gain, HIV, adipogenesis, 3T3-L1 cells, cytomatrix, integrase strand transfer inhibitors, nucleoside reverse transcriptase inhibitors

## Abstract

Integrase strand transfer inhibitors (INSTI) are a recently available class of antiretroviral therapy (ART) medications with a good tolerability profile and a high genetic barrier to HIV drug resistance. However, several studies report more significant weight gain among persons receiving INSTI-based ART regimens for initial therapy compared to protease inhibitors (PIs) and nucleoside reverse transcriptase inhibitors (NNRTI)-based regimens. In our experimental setting, we used the *in vitro* model of adipogenesis of 3T3-L1 cells to investigate the effects of the NRTIs tenofovir disoproxil fumarate (TDF) and tenofovir alafenamide (TAF), alone or in combination with four integrase strand transfer inhibitors: raltegravir (RAL), elvitegravir (ELV), dolutegravir (DTG) and bictegravir (BIC) on adipose differentiation. In addition, protein expression levels of PPARɣ and C/EBPα, and the intracellular lipid accumulation by Red Oil staining, were used to monitor adipocyte differentiation. Compared to control, RAL, ELV, DTG, and BIC were all able to increase adipogenesis, being in this, RAL and ELV more efficient. On the other hand, TAF and TDF inhibited adipogenesis. Moreover, when used in combination with the other INSTI molecules, TAF and TDF were able to reduce the adipogenic effects of all four drugs. This ability was more evident when TAF was used in combination with DTG and BIC. All these data suggest that TAF and TDF have an inhibitory effect on adipogenesis *in vitro* and that they could also effectively counteract the increased adipogenesis caused by the treatment with INSTIs. Finally, to evaluate if the 3T3-L1 cell could express fibroblast-like features following INSTIs treatment, we evaluated the immunohistochemical expression of ER-TR7, a well-known fibroblastic marker. This last assay showed that treatment with INSTIs increased the expression of ER-TR7 compared to control and to cells treated with TAF o TDF.

In conclusion, our experimental data support the evidence that *in vitro* challenge of 3T3-L1 cells with INSTIs is able to increase adipocytic differentiation and to drive a number of these cells toward the expression of fibroblastic features, with a different degree according to the various drugs used, while TAF and TDF have an antagonistic role on this phenomenon.

## 1. INTRODUCTION

Human immunodeficiency virus (HIV) infection was previously related to a high prevalence of wasting, being this a negative prognostic indicator of HIV disease progression, nutritional depletion, and susceptibility to opportunistic infection [1].

Modern antiretroviral therapy (ART) currently allows people living with HIV (PLWH) to survive decades on treatment because of its excellent antiviral efficacy and tolerability, proven durability against resistance mutations together to the ability to reverse the catabolic state and reduce circulating inflammatory markers, to improve appetite and nutrient absorption [2] and to reduce the risk of wasting [3].

However, this clinical success has been accompanied by an increasing proportion of overweight and obese PLWH, in both resource-poor and resource-rich environments, with women being more affected than men [3, 4].

Obesity is associated with adipose tissue dysfunction and inflammation, increasing the aisk for metabolic diseases, including diabetes mellitus, neurocognitive impairment, liver disease, and cardiovascular disease [5]. In addition, weight gain in PLWH confers a greater risk of metabolic disease than in HIV-negative individuals [6].

The mechanisms contributing to weight gain on ART are not entirely understood and could reflect off-target effects of mechanisms or greater efficacy and suppression of viral reservoirs that reduce the metabolic cost of HIV infection.

Both HIV infection and ART target the adipose tissue. Several factors could explain the role of the infection in weight gains, such as mechanisms relating to adipose dysfunction and fibrosis, immune function, inflammation, and gastrointestinal loss of integrity. In addition, the return-to-health effect in individuals effectively treated with ART could also play a role [7].

Many studies reported the role of ART medications in weight gain among PLWH. Overall, no convincing differential effect of the nucleoside reverse transcriptase inhibitors (NRTIs) or non-nucleoside reverse transcriptase inhibitors (NNRTIs) classes has been observed. The older NNRTIs such as Efavirenz (EFV) and NRTIs, such as the thymidine analogues (zidovudine and stavudine), lamivudine, abacavir, and tenofovir disoproxil fumarate (TDF), predominantly target mitochondrial DNA, inducing oxidative stress and adipocyte death, but they have not been associated with differential weight gain [8, 9]. The role of protease inhibitors (PIs) also remains uncertain, but PIs do not generally lead to more total or central obesity than NNRTIs [10, 11]. Integrase strand transfer inhibitors (INSTIs) are a more recently available class of ART medications with a good tolerability profile and genetic barrier to HIV drug resistance. This new class of drugs currently includes raltegravir (RAL), elvitegravir (ELV), dolutegravir (DTG), and bictegravir (BIC) [12, 13]. INSTI-based ART regimens are now recommended as first-line treatment for most PLWH [14], but several recent studies, generally from single sites or cohorts, report more significant weight gain among persons receiving INSTI-based ART regimens for initial therapy as compared to PIs and NNRTI-based regimens. The first developed INSTI, raltegravir (RAL), led to similar visceral adipose tissue accumulation when compared to PIs [15] in randomized studies, but it was associated with more considerable gains in waist circumference and a higher incidence of severe weight gain through 96 weeks of ART [16, 17]. In a cohort from Brazil, PLWH on RAL-based regimens were 7-fold more likely to become obese compared to those receiving NNRTI- or PI-based regimens [18]. Moreover, observational and cohort studies suggest that INSTIs can lead to excessive gains in body weight, being, in this, DTG and RAL more efficient than ELV [18-21]. In other observational studies, INSTI-based regimens, particularly DTG-based ART regimens, were associated with greater weight gain [19, 21-23].

In vitro studies demonstrated that DTG inhibited the binding of the radiolabelled α-melanocyte-stimulating hormone to the human recombinant melanocortin-4 receptor. This phenomenon may interfere with the regulation of food intake and lead to obesity [24]. However, McMahon et al. subsequently found that drug concentrations substantially greater than clinical exposure are required for antagonism of the melanocortin-4 receptor to occur, thus refuting this hypothesis [25].

To date, the use of the newer NRTI, tenofovir alafenamide fumarate (TAF), is rapidly increasing, given the lower incidence of adverse bone and renal effects compared to TDF [26, 27]. In addition, recent data from randomized trials and smaller observational studies indicate that TAF use could predispose to weight gain in association with or independently of concomitant INSTI use [28, 29].

Adipogenesis represents an example of terminal differentiation cells with the cessation of cell proliferation, the accumulation of cells in the G1 phase of the cell cycle, and, later on, the activation of specific transcription factors [30]. Indeed, adipocyte differentiation is largely controlled by two families of transcription factors: the CCAAT/enhancer-binding proteins (C/EBPs) and the peroxisome proliferator-activated receptors (PPARs). In the early phase of adipocyte differentiation, the different members of the C/EBP family form heterodimers among them and are able to bind and regulate, through a DNA binding domain, the expression of many genes fundamental for the early steps of adipogenesis. Instead, in the second phase of adipocyte differentiation, the different members of the PPARs family form heterodimers among them and are able to bind and activate specific nuclear hormone receptors representative of the mature adipocyte [31]. Interestingly, the evaluation of the expression of adipocytic differentiation markers, such as PPAR-γ and C/EBP-α, confirmed the impairment of fat tissue differentiation in HIV patients [32].

Many studies also suggested that disturbances in adipose tissue gene expression are already present in untreated HIV-1 infected patients, thus indicating a role of HIV-1 itself in eliciting adipose tissue alterations that could be worsened by HAART, which ultimately leads to HALS (HIV-associated lipodystrophy) [33].

Furthermore, in the last 20 years, some studies reported increased fibrosis in adipose tissue from HIV-infected patients [34-36], and several mechanisms have been hypothesized to explain this phenomenon. It has been described the increase of extracellular matrix in adipose tissue by deposition of collagen produced by adipocytes [35, 37] and the reprogramming of adipocytes into fibroblast, confirmed by the expression of ER-TR7, a well-known fibroblastic marker was also reported [38]. In addition, more recently, Ngono et al. described the ability of BIC and DTG to induce a pro-fibrotic phenotype in simian beige adipose tissue and found this phenotype to be associated with insulin resistance [39].

Drawing from this background, we decided to evaluate the effects of the treatment with four integrase strand transfer inhibitors (ELV, RAL, DGT, and BIC) alone or in combination with TAF or TDF on 3T3-L1 cells during adipose differentiation.

## 2. MATERIALS AND METHODS

### 2.1 CELL CULTURE

3T3-L1 preadipocytes were cultured and differentiated using a standard protocol [40, 41]. 3T3-L1 cells were cultured in Dulbecco’s modified Eagle’s medium supplemented with 10% fetal bovine serum in a humidified atmosphere of 5% CO_2_ 95% air at 37°C until confluence. Two days later, induction of adipocyte differentiation was initiated by treatment of cells with a differentiation medium containing 1 mM insulin, 1 mM Dexamethasone, and 0.5 mM isobutylmethylxanthine for 2 days, followed by 2 days of treatment with a medium containing 1 mM insulin alone. The medium was replaced every 2 days for the following 8 days. Treatment with drugs (TAF, TDF, ELV, RAL, DTG, BIC) was started on the first day of induction and added daily until day 4 of differentiation. All drugs were used in single and combination with TAF and TDF, all at a concentration of 30 μg/ml. All drugs were obtained from Vinci-Biochem, IT.

### 2.2 CELL VIABILITY ASSAY

The effects of the drugs on cell viability were determined by the MTT assay as previously described with some modifications [42]. In brief, cells (10,000 cells per well) were incubated with or without the drugs in three different experiments in a 96-well plate and incubated for 24-48 h at 37 °C. Our control is given by cycling 3T3-L1 cells. After incubation, 10 μl of MTT (3 [4,5 dimethylthiazol 2yl] 2,5 di-phenyl-tetrazolium bromide, GoldBio.Com) solution (5 mg/ml) were added to each well and incubated for another 4 h at 37°C. The supernatants were aspirated carefully, and 100 μl of DMSO were added; then, the plate was held on a vibrator for 20 s. The optical density of the cell suspension was measured at 570 nm using a microplate reader (Bio-Tek Instruments Inc., Winooski, VT). Cell viability was expressed as a percentage of MTT reduction, assuming that the absorbance of untreated cells was 100%.

### 2.3 PROTEIN EXTRACTION AND WESTERN BLOTTING ANALYSIS

3T3-L1 cells were lysed in Ripa buffer for 15 minutes on ice. Total extracts were cleaned by centrifugation for 15 minutes at 4°C at 10.000 rpm and analyzed for protein content by the Bradford method. Twenty micrograms of protein from each cell lysate were separated by 12% SDS-PAGE and transferred to PVDF membranes, and the filters were stained with 10% Ponceau S solution for 2 minutes to verify equal loading and transfer efficiency. The membranes were tested with primary antibodies anti-PPARɣ (1:200) and anti-C/EBPα (1:200) (Santa Cruz Biotechnology, CA) and incubated overnight. Then the membranes were incubated with 1:5000 HPR-conjugated anti-mouse and anti-rabbit antibodies for 1 h at room temperature. They were extensively washed and finally analyzed using the ECL system (Amersham). Anti-β-tubulin (Elabscience Biotechnology Inc., US) was used to estimate the equal protein load. The experiments were repeated three times, obtaining comparable results.

Imagej software was used to relatively quantify protein expression levels from western blot analysis.

### 2.4 RED OIL O STAINING

Intracellular lipid accumulation was determined by Red Oil O staining (Sigma-Aldrich®) during adipocyte differentiation. Cells were washed twice with ice-cold buffered saline (PBS), fixed with 4% (w/v) paraformaldehyde in PBS for 20 min, and stained with a 0.5% (w/v) red oil O solution in isopropanol for 30 min at room temperature. After staining, cells were washed with PBS to remove the excess stain before photography.

### 2.5 IMMUNOHISTOCHEMISTRY

Adipogenesis was induced in 3T3-L1 cells alone or treated with TAF, TDF, ELV, RAL, DTG, and BIC. On day 8 of differentiation, cells were recovered. Then, 2×10^5^ cells/ml cells were loaded into Cytomatrix (UCS Diagnostic S.r.l., Rome, Italy) and entrapped in this sponge by formalin fixation, as already described [43-45]. After fixation, Cytomatrix-entrapped cells were included in paraffin following standard protocols, and sections were cut at 5 µm using a microtome LEICA SM 2000R (Advanced Research System Inc., Macungie, PA). Sections were dewaxed in xylene, rehydrated through a series of graded ethanol solutions, and stained with Gill’s Haematoxylin and Eosin (Bio-Optica, Milan). Immunohistochemistry was executed on consecutive sections using the anti-fibroblast monoclonal antibody ER-TR7 (Invitrogen MA1-40076). Tissue sections were sequentially treated with 3% hydrogen peroxide in an aqueous solution and blocked with 6% milk in PBS. The slides were then incubated for one hour at room temperature with the ER-TR7 antibody, at a final dilution of 1:100. Following three PBS washes, slides were incubated with UltraTek HRP secondary antibody (ScyTek Laboratories, Logan, Utah, U.S.A.) for 1 hr at room temperature Diamonibenzidine (ScyTek Laboratories, Logan, Utah, U.S.A) was used as the final chromogen and hematoxylin was used as the nuclear counterstain. Each tissue section’s negative control was generated without the primary antibody. All samples were processed under the same conditions. The cellular expression levels of ER-TR7 per field (20X) were calculated under the microscope by two different observers (ADL and AB) and characterized as follows: score 0 (absent), score 1 (low; up to 5% of positive cells), score 2 (moderate; up to 10% of positive cells), score 3 (high; more than 10% of positive cells). An average of 10 fields was observed for each sample. Concordance among the observers was found in all the samples analyzed.

### 2.6 STATISTICAL ANALYSIS

We used Excel for statistical analysis. Where applicable, results were expressed as the mean ± standard deviation (SD) of at least three independent experiments. Statistical differences were considered if the p-value was < 0.05*, p-value was < 0.01**, p-value was < 0.001*** as determined by ANOVA followed by Student’s t-test.

## 3. RESULTS

3T3-L1 cells were induced to differentiation, adding at time 0 the nucleotide analogues TAF and TDF in single and in combinations with four integrase inhibitors (ELV, RAL, DGT, and BIC). The control point was obtained by inducing 3T3-L1 cells to differentiation without using the drugs. We performed an MTT assay to define the concentration not toxic for the treatment. All the drugs were used at different concentrations from 15 to 50 μg/ml (15-20-30-50 μg/ml) for 24h (Figure 1A) and 48h (Figure 1B). No reduction in cell viability was observed at any point, and the concentration chosen for all subsequent experiments was 30 μg/ml for all drugs (figure 1 A, B).

**Figure 1:**
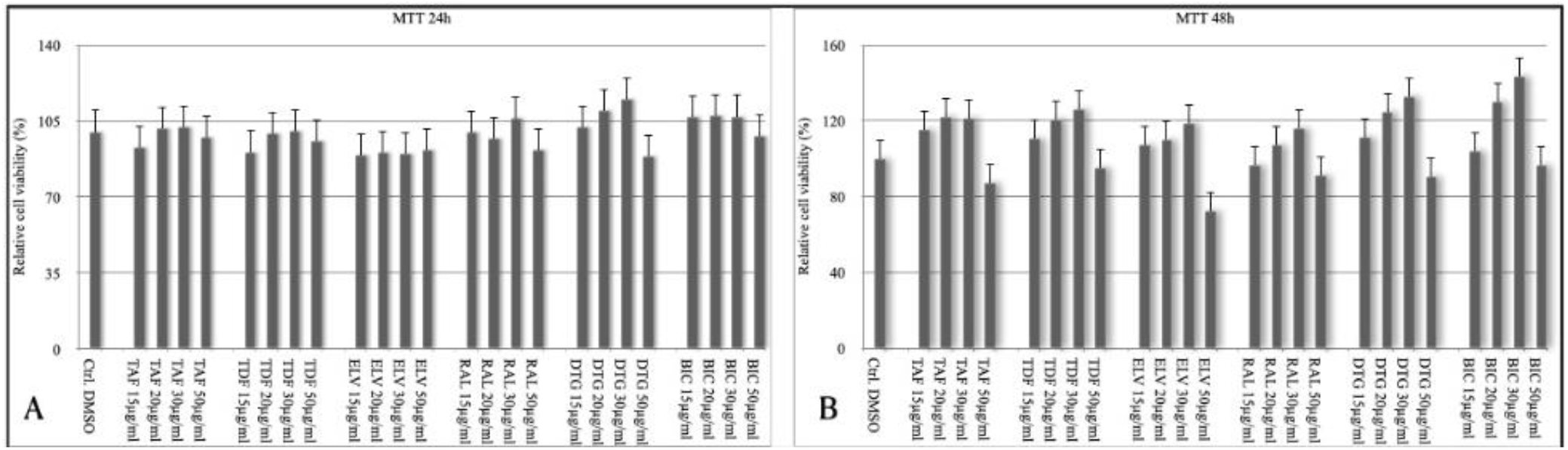
MTT assay. All the drugs were used at different concentrations from 15 to 50μg/ml (15-20-30-50μg/ml) for 24h (A) and 48h (B). No reduction in cell viability was observed at any point.

3T3-L1 cells were treated during adipose differentiation by adding the nucleotide analogues TAF and TDF. The treatment with TAF was able to induce a slight reduction in the number of lipid droplets stained by Red Oil in the cytoplasm compared with the untreated cells (Figure 2A). Furthermore, when looking at the expression levels of PPARɣ and C/EBPα, an up-regulation of these is observed, moving from the first to the third point of differentiation, but to a reduced extent compared to the control curve of the non-pharmacologically treated cells (Figure 3 A, B, C).

**Figure 2:**
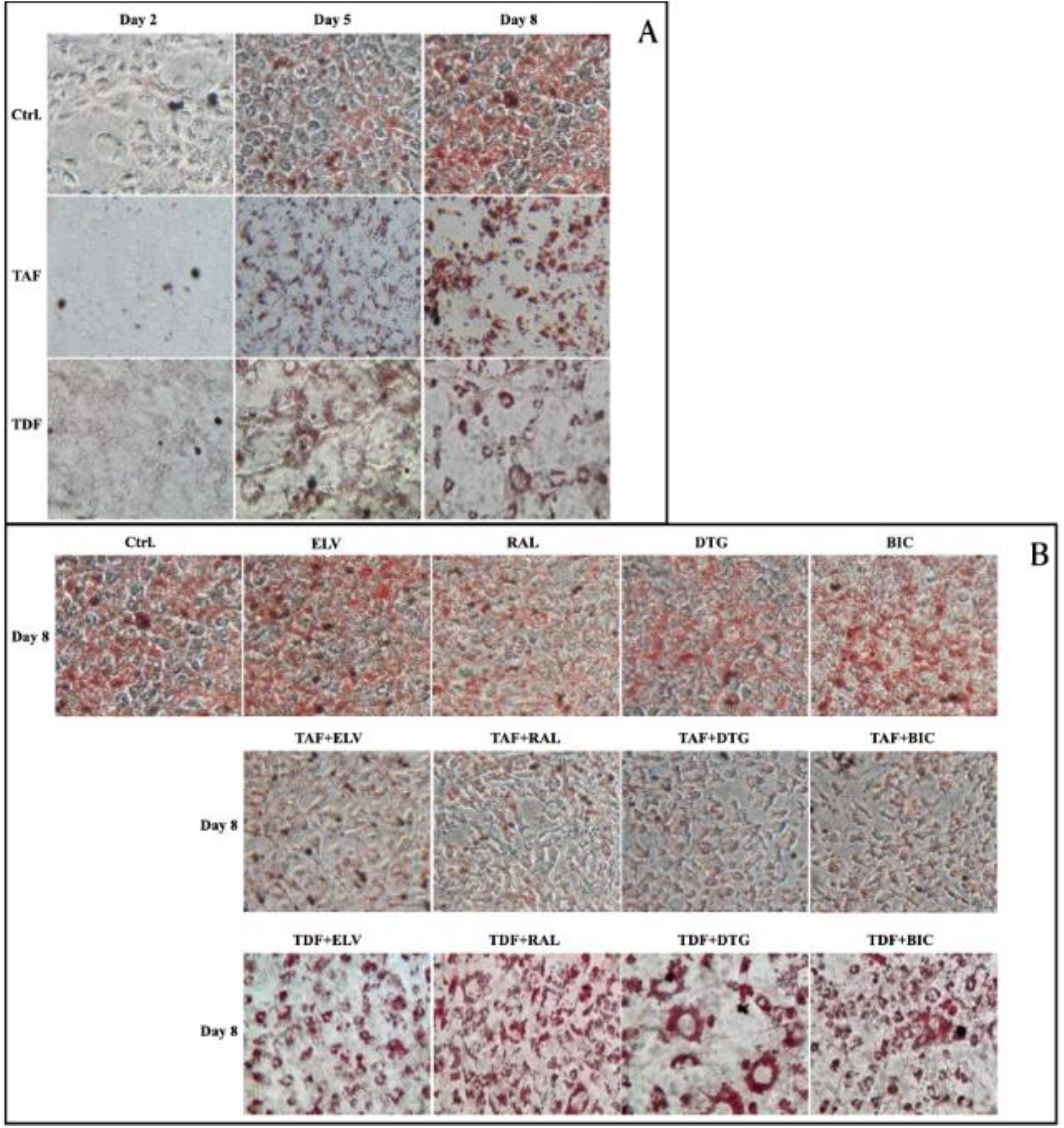
Red Oil O staining. (A) The treatment with TAF and TDF was able to induce a slight reduction in the amount of lipid droplets in the cytoplasm compared with the untreated cells; concerning the morphology, cells appeared smaller and more elongated with respect to the control. (B) Treatment with the four integrase inhibitors (ELV, RAL, DGT and BIC) was able to differentiate 3T3-L1 into adipocytes, as can be seen by the increase in the amount of lipid droplets on day 8 of differentiation, compared to the control. Treatment with TAF or TDF in combination with the four integrase inhibitors (ELV, RAL, DGT and BIC) showed a decrease in the amount of lipid droplets in the cytoplasm compared to untreated cells. Concerning morphology, a phenotype similar to the one observed in cells treated with TAF and TDF alone, was found.

**Figure 3:**
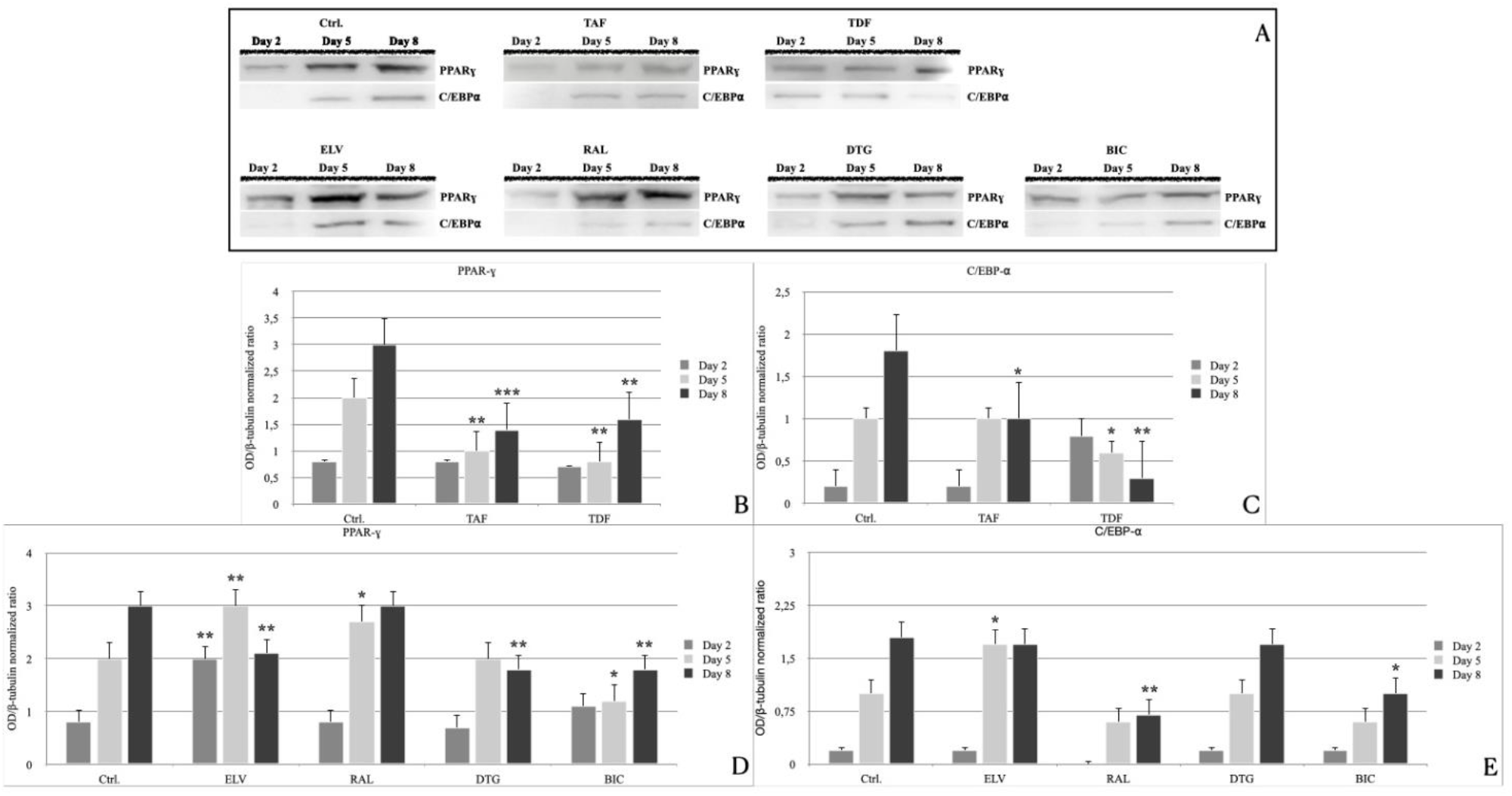
Western Blotting. Treatment with TAF was able to induce a constant decrease in protein expression of PPARɣ and C/EBPα, from the first to the third point of differentiation, compared to control(A-C); Treatment with TDF was able to decrease the protein expression of PPARɣ and C/EBPα, from the first to the third point of differentiation compared to control(A-C); Treatment with ELV and RAL was able to increase the expression levels of PPARɣ since day 5, compared with the control (A, D), while at day 8 the expression of PPARɣ was still higher in cells treated with RAL, while ELV treatment displayed a decrease of the expression levels of this protein (A, D). The expression levels of C/EBPα for both these drugs were slightly lower than in the untreated cells (A, E); Treatment with DTG and BIC was able to increase the expression of PPARɣ and C/EBPα between day 2 and day 5 slightly and was constant between day 5 and 8, compared with the control (A, D, E). All data were normalized againstβ-tubulin obtained on the same immunoblotting membrane after stripping. The resulting ratio of the target protein to tubulin was used for statistical analysis and means ±SD are shown as bar graphs. p < 0.05*, p < 0.01**, p< 0.001***

Treatment with TDF caused a reduction in the number of lipid droplets stained with Red Oil in the cytoplasm compared to the untreated cells and an overall reduced number of cells (figure 2A). Consistently, even if an up-regulation of PPARɣ from the first to the third point of differentiation was observed, it was reduced compared to the control, whereas a C/EBPα down-regulation was observed at the third point of differentiation (figure 3 A, B, C). Indeed, in both treatments, cell morphology did not reflect the classic adipose cell features characterized by a rounded shape with the droplets arranged around the nucleus but appeared smaller in size and more elongated (figure 2A). Moreover, we observed also a reduction in the total number of cells, suggesting a cell proliferation slowdown which was much more evident for TDF.

3T3-L1 cells during adipose differentiation were then treated by adding independently four integrase inhibitors (ELV, RAL, DGT, and BIC). The four drugs were able to differentiate the 3T3-L1 to adipocyte, as demonstrated by the amount of lipid droplets stained with Red Oil at day 8 of differentiation, which was higher when compared to the control (figure 2A). More in detail, RAL and ELV were able to increase the number of lipid droplets in the cytoplasm more than DGT and BIC (figure 2B). Concerning morphology, cells were very similar to the control in number and shape.

When we looked at the protein expression of PPARɣ and C/EBPα, we found that, compared with the control, cells treated with ELV and RAL were able to increase the expression levels of PPARɣ since day 5. On day 8, the expression of PPARɣ was still higher in cells treated with RAL, while ELV treatment displayed a decrease in the expression levels of this protein (Figure 3A-D). The expression levels of C/EBPα for both these drugs were slightly lower than in the untreated cells being RAL even more effective in this effect (figure 3 A, E). In the cells treated with DTG and BIC, the expression of PPARɣ and C/EBPα increased slightly between day 2 and day 5 and was constant between day 5 and 8 (Figure 3 A, D, E).

When TAF was added in combination with the four integrase inhibitors (ELV, RAL, DGT, and BIC), we observed a decrease in the amount of lipid droplets in the cytoplasm compared with the untreated cells (Figure 2B). The ability of TAF to inhibit the differentiation of 3T3-L1 cells was more evident in combination with DTG and BIC (Figure 2B). Coherently, the differentiation treatment with TAF in combination with the four integrase inhibitors induced a decrease in the expression levels of PPARɣ from day 5 to 8. In particular, we observed that PPARɣ at day 8 was still expressed in cells treated with TAF in combination with ELV or RAL, while PPARɣ expression was almost absent in cells treated with TAF in combination with DTG or BIC (Figure 4B, D). Finally, in all the combinations, the expression levels of C/EBPα were slightly higher than those obtained in the treatment of cells with the four integrase inhibitors without TAF (Figure 4B, E).

**Figure 4:**
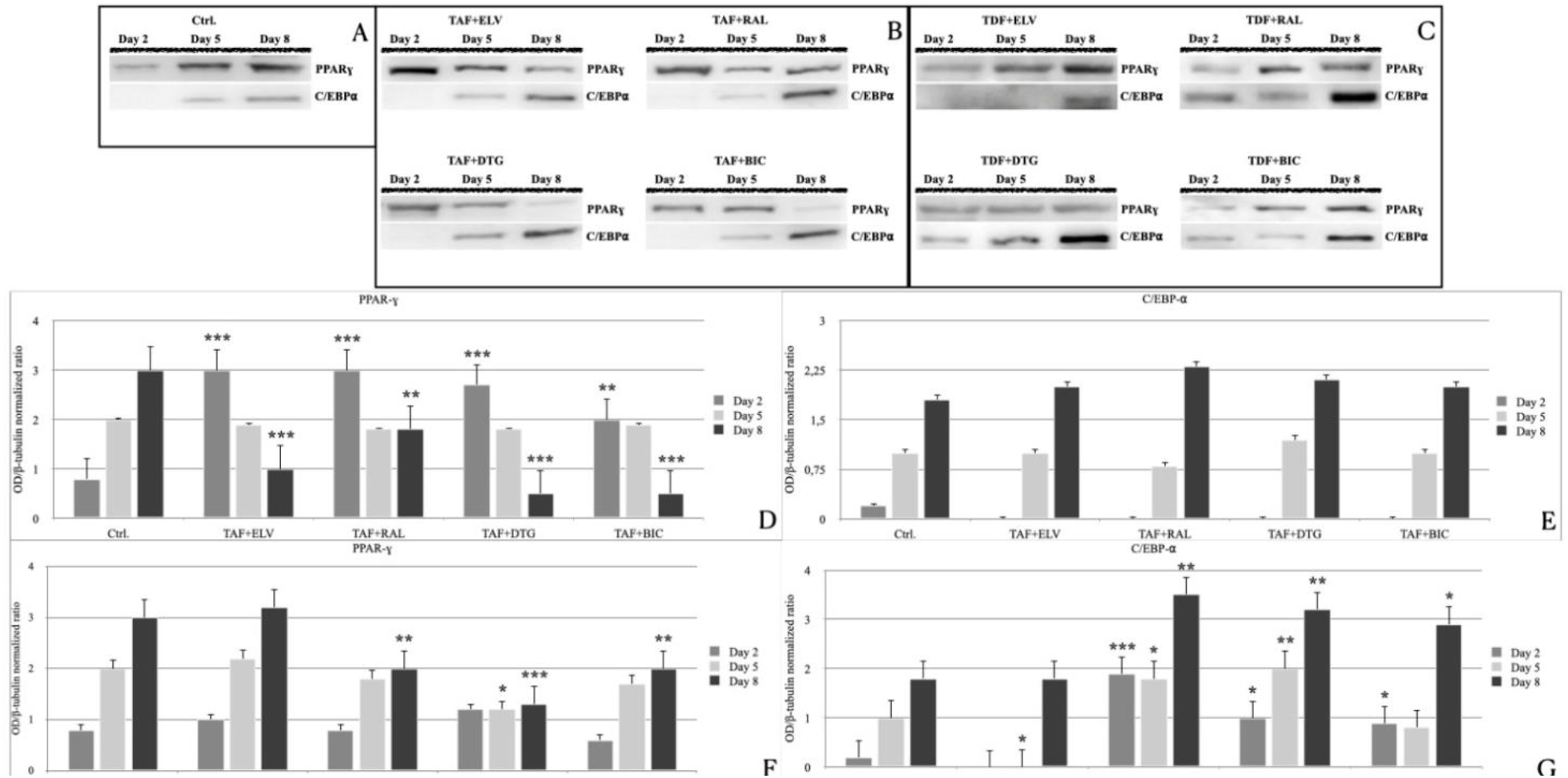
Western Blotting. Treatment with TAF and INSTIs induced a decrease in the expression levels of PPARɣ from day 5 to 8. PPARɣ at day 8 was still expressed in cells treated with TAF in combination with ELV or RAL, while in cells treated with TAF in combination with DTG or BIC it was almost absent (B,D); in all the combinations the expression levels of C/EBPα was similar to the expression obtained in the treatment of cells with the INSTI without TAF (B, E); Treatment with TDF was able to induce steady decrease in the expression of PPARɣ (C, F). An increase in protein level for C/EBPα, from the first to the third point of differentiation, was observed (C, G). All data were normalized againstβ-tubulin obtained on the same immunoblotting membrane after stripping. The resulting ratio of the target protein to tubulin was used for statistical analysis and means ±SD are shown as bar graphs. p < 0.05*, p < 0.01**, p< 0.001***

When we looked at the TDF treatment of 3T3-L1 cells in combination with the four INSTIs, we found that this last drug was still able to induce differentiation. Furthermore, by means of Red Oil staining, intense agglomerates of lipid droplets were observed in the cytoplasm of cells.

Treatment with TDF was able to induce up-regulation of both PPARɣ and C/EBPα in all combinations from day 2 to day 8, especially comparing the points with TAF combined with the four integrase inhibitors, and in particular of C/EBPα, in the third point, in combinations with DTG and RAL (figure 4C, F, G). Challenging preadipocytes with TDF plus INSTIs determined an up-regulation of both PPARɣ and C/EBPαfrom the first to the third point of the differentiation. This means that in all combinations, but the one with ELV, PPARɣ was less up-regulated concerning the control cells, while C/EBPa, was more expressed at the last point, with respect to the controls and the TAF plus INSTs combinations. This behaviour was more evident with RAL and DTG. Therefore, TDF plus INSTIs was related to an increase in induction to differentiation compared to control and even more evident in comparison with TAF plus INSTIs administration (figure 4C, F, G). Concerning the morphology, cells appeared smaller and more elongated with respect to the control (Figure 2B).

Finally, both TAF and TDF, when used in combination whit the various INSTIs, were able to cause a decrease in the number of cells, being TDF much more effective. Interestingly, these effects on the phenotype of the cells, both in terms of number and shape, caused by treatment with TAF and TDF alone, were also detected when these drugs were used in combination with the four INSTIs, being TDF more effective.

Following the observed morphological changes, to assess whether these cells could be somehow redirected towards a fibroblast-like phenotype at certain points of adipogenesis, we evaluated the expression of ER-TR7, a known fibroblast marker in 3T3-L1 cells treated with ART.

Table 1 summarizes the data obtained with immunohistochemistry for ER-TR7 expression in 3T3-L1 cells after 8 days of induction of adipogenesis. ER-TR7 expression was detected in all the different treated 3T3-L1 cells. In detail, expression of ER-TR7 was low in control cells, and a comparable result was also obtained in cells treated with TAF and TDF. Interestingly, cells treated with BIC and RAL displayed a moderate expression for ER-TR7, while cells treated with ELV and DTG exhibited a high expression of ER-TR7 (Figure 5).

**Table 1:** Summary of data obtained with immunohistochemistry for ER-TR7 expression in 3T3L-1 cells after 8 days of induction of adipogenesis.

| Drug | TR7 immunohistochemical score |
| --- | --- |
| Ctrl | Low |
| TAF | Low |
| TDF | Low |
| ELV | High |
| RAL | Moderate |
| DTG | High |
| BIC | Moderate |

**Figure 5:**
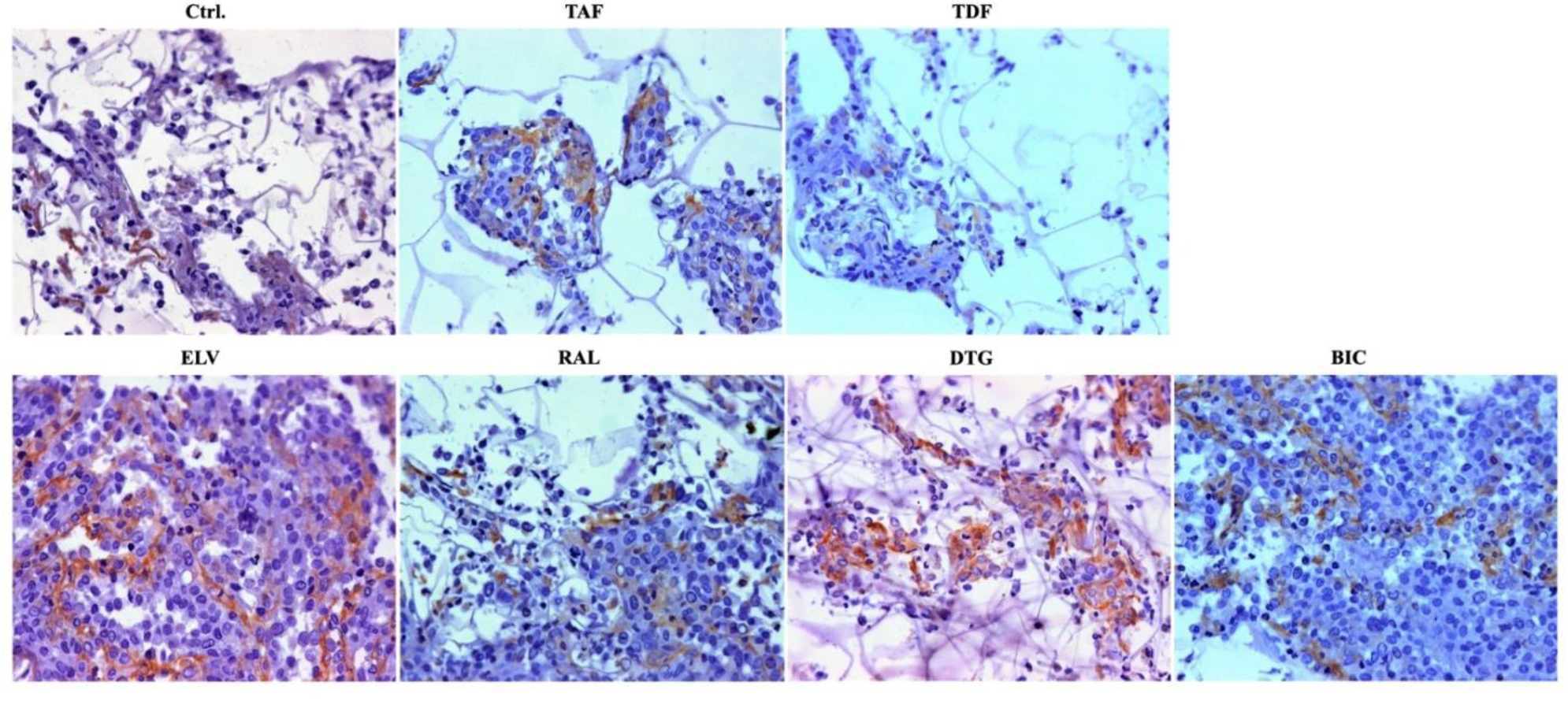
Immunohistochemistry. Expression of TR7 was low in control cells, and a comparable result was also obtained in cells treated with TAF and TDF. Treatment with BIC and RAL displayed a moderate expression for TR7, while cells treated with ELV and DTG exhibited a high expression of TR7.

## 4. DISCUSSION

Adipose fat tissue contributes significantly to total body weight ranging from approximately 20-30 % in non-obese individuals to 50% in persons with obesity, white adipose tissue represents the significant components of fat mass compared to brown and beige fat [46]. White adipocytes are universally recognized as a storage compartment of dietary lipids for further use during fasting periods. However, the role of adipose tissue is more complex, displaying relevant metabolic activities as well as well-known roles in inflammation, endocrine-immune-function, and tissue repair [34, 35]. In people living with HIV (PLWH), adipose tissue is targeted by both the virus and antiretroviral drugs. Therefore, it is challenging to distinguish the different contributions of these two factors to adipose tissue altered functions in PLWH [6, 47].

Drawing from this background, we decided to assess in the in vitro model of adipogenesis of 3T3-L1 cells the contribution of antiretroviral drugs to adipocyte differentiation potential impairment. In addition, considering some observations reporting the increase of fibrosis in adipose tissues in PLWH [34-36], we further investigated the expression of a specific fibroblast marker ER-TR7 under pharmacological challenge. Our *in vitro* results confirm that INSTIs could induce adipogenesis with a different degree of efficacy according to the drug used, consistently with clinical and experimental observations [15-23, 48]. This provides an alternative hypothesis for inhibiting the binding of the α-melanocyte stimulating hormone to the human recombinant melanocortin 4 receptor, described in vitro for DGT [25]. In our experimental setting, DGT and ELV were more efficient in the induction of adipogenesis. Several latest published data seem to confirm our observation. In a recent report, Gorwood et al. [36] showed the ability of DTG and RAL to exert pro-adipogenic effects in human/simian adipose tissue and human adipocytes, and Ngono et al. confirmed this result in an animal model of SIV-infected macaques, where DTG was associated with higher adipogenesis [39]. In addition, we found that TAF treatment had an inhibitory effect on adipocytic differentiation, acting in the earlier phases through PPARγ expression deregulation.

Interestingly, this action of TAF was able to lead to an antagonistic effect on adipocyte differentiation and effectively counteract the increase in adipogenesis caused by INSTIs, in particular DTG and BIC. This evidence seems in sharp contrast with clinical studies indicating that TAF could determine weight gain [28, 29]. However, when analyzing these data in an in vivo setting, we must consider the multifactorial nature of this phenomenon. On the other hand, TDF displayed a similar effect on adipogenesis both when used alone or in combination with INSTIs. More in detail, it seems to interfere with adipogenesis mostly at the last step of the differentiation, affecting C/EBPα expression levels and having as a result a more marked induction of the differentiation. This effect of TDF in adipogenesis needs to be further elucidated with more detailed evidences from experimental and clinical settings also to better define this phenomenon in accordance with the well-known “statin-like effect” of this drug in clinical settings [49]. Interestingly, we noticed that TAF and TDF exerted an inhibition in cell proliferation and an effect on cell morphology, being TDF more efficient than TAF. This phenomenon was evident when the drugs were used alone, but also when used in combination with the INSTIs molecules. On the other hand, the INSTIs drugs did not display this ability, at least in our experimental setting.

When we looked at the expression of fibroblastic features in our terminally differentiated 3T3-L1 cells, we found that INSTIs were all able to increase the expression of ET-R7 compared to TAF, TDF, and the control. DTG and EVT were more efficient in this activity. This last result acquires high relevance considering recent reports showing that forced expression of PPARɣ is able and sufficient to induce adipogenesis in fibroblasts [36] and that adipocytes can be reprogrammed into fibroblasts under transfection of some master regulators [38]. More recently, Jones et al. demonstrated that adipocytes could acquire a fibroblast-like transcriptional signature by using a murine animal model under over-feeding conditions [50]. Considering all this evidence together with our experimental data, one can support the hypothesis of a partial reconversion from adipocytes into fibroblastic phenotype under specific conditions in our experimental setting.

We can conclude that our *in vitro* study confirms that INSTIs could induce adipogenesis and increase the pro-fibrotic activity of adipocytes, while interaction between INSTIs and TAF or TDF leads to an antagonistic effect on the differentiation of adipocytes.

The mechanisms of TAF and TDF in modifying adipogenesis, as well as the ability of INSTs to induce fibrosis production in adipose tissue, need to be further elucidated with more detailed evidence from experimental and clinical settings.

